# Neural Correlates of Reactivation and Vividness Reveal Separable Contributions to Objective and Subjective Measures of Episodic Memory

**DOI:** 10.1101/2021.03.11.434635

**Authors:** Ryan M. Barker, Marie St-Laurent, Bradley R. Buchsbaum

**Affiliations:** Department of Psychology, University of Toronto; Rotman Research Institute, Baycrest Hospital; Université de Montréal et Mila

**Keywords:** episodic memory, vividness, neural reactivation, meta-analysis, hippocampus

## Abstract

Episodic recollections vary in fidelity, sharpness, and strength—qualities that can be examined using both introspective judgements of mental states and objective measures of brain activity. Subjective and objective measures are both valid ways of “reading out” the content of someone’s internal mnemonic states, each with different strengths and weaknesses. St-Laurent and colleagues (2015) investigated the neural correlates of memory vividness ratings and neural reactivation during memory recall and found considerable overlap, suggesting common neural basis underlying these different markers of successful recollection. Here we extended this work with a much more extensive examination in which we used meta-analytic methods to pool four neuroimaging datasets in order to compare and contrast the neural substrates of neural reactivation and vividness judgements. While reactivation and vividness judgements correlated positively with one another and were associated with common univariate activity in the dorsal attention network and anterior hippocampus, some differences were also observed. Vividness judgments were tied to stronger activation in the striatum and dorsal attention network, together with suppression of default mode network nodes, and we also observed a trend for reactivation to be more closely associated with early visual cortex activity. A mediation analysis found support for the hypothesis that neural reactivation is necessary for vivid recollection, with activity in the anterior hippocampus associated with greater reactivation. Our results suggest that neural reactivation and vividness judgements reflect common recollective processing but differ in the extent to which they engage effortful, attentional processing. Additionally, the similarity between reactivation and vividness appears to arise, partly, through hippocampal processing during recollection.

## 1. Introduction

Recollection is the subjective experience of projecting oneself mentally back in time to re-experience an event from one’s past. Not all recollections, however, are alike. Some recollections evoke rich perceptual detail, as if one were reliving the moment again. Other recollections may be less rich in detail, with the recollected context emerging as more vague or fuzzy (i.e., recollection has been shown to vary along dimensions of precision: Harlow & Yonelinas, 2014; Richter, Cooper, Bays, Simons, 2016; confidence: Ingram, Mickes, & Wixted, 2012; and amount of information represented: Vilberg, Moosavi, & Rugg, 2006; Vilberg & Rugg 2007; Vilberg & Rugg, 2009), and there are gradations in the phenomenal experience accompanying recollection that lie in between these extremes. A memory researcher has no easy way to directly measure the “quality” of recollection. An introspecting human subject, however, is capable of reporting and rating various aspects of their conscious experiences. A word in the English language often used to describe the *sharpness* and vitality of a conscious experience is “vivid” or “vividness”. Psychologists often ask participants in experiments to rate their conscious memories (or mental images) as having more less of this quality (e.g., Cooper, Kesinger, & Ritchey, 2019; Houben, Otgaar, Roelofs, Merckelbach, & Muris, 2020; Janssen, Rubin, & St. Jacques, 2011; Kark & Kesinger, 2019; Tibon, Fuhrmann, Levy, Simons, & Henson, 2019). In practice, “vividness” provides an index of a memory’s subjective global quality and the extent to which a recalled memory has something of the force and sharpness of a direct perceptual experience.

The phenomenology of successful recollection seems to mimic perception itself, and the experiential similarity between the perception of an event and its recreation in the mind suggests that certain underlying brain processes are shared between both experiences (Alvarez & Squire, 1994; Buchsbaum, Lemire-Rodger, Fang, & Abdi, 2012; Damasio, 1989; Hebb, 1968). Indeed, accurate memory for source features has been associated with reactivation of brain states recruited during the occurrence of the previously encoded event (e.g., Gordon, Rissman, Kiani, & Wagner, 2014; Kuhl, Johnson, & Chun 2013; Ritchey, Wing, LaBar, & Cabeza, 2013; Staresina, Henson, Kriegeskorte, & Alink, 2012), with stronger correlations when participants report greater confidence (Kim, Norman, & Turk-Browne, 2019; Thakral, Wang, & Rugg, 2015) and specificity (Kuhl, Rissman, Chun, & Wagner, 2011) of source memory judgements, as well as freely-recalled auditory and visual features (St-Laurent, Abdi, Bondad, & Buchsbaum, 2014). Therefore, like vividness judgements, brain reactivation measures capture the content and quality of memory, however this approach offers a purely quantitative, objective metric.

Previous work has established a clear relationship between subjective vividness judgements and objective neural reactivation measures in the context of recollection on a trial-by-trial basis (e.g., Bird, Keidel, Ing, Horner, & Burgess, 2015; Bone et al., 2019; Johnson, Kuhl, Mitchell, Ankudowich, & Durbin, 2015; St-Laurent, Abdi, and Buchsbaum, 2015). Studies examining the shared neural substrates underlying vividness and neural reactivation have shown a correlation between these two measures in posterior sites tied to perceptual processing, reflecting the importance of perceptual activity for vivid remembering and suggesting that reactivated patterns may provide a neural context from which vivid memories emerge (Bone et al., 2019; Bone, Ahmad, & Buchsbaum, 2020; Buchsbaum et al., 2012; Dijkstra, Bosch, & van Gerven, 2017; St-Laurent et al., 2015; Wing, Ritchey, & Cabeza, 2015).

St-Laurent, Abdi, and Buchsbaum (2015) used a data-driven approach to identify brain regions conjointly associated with neural reactivation and vividness, irrespective of the specific content of recollected episodes. Vividness ratings and whole-brain neural reactivation, measured as the similarity between distributed brain activity patterns evoked by the encoding and retrieval of short multimodal video clips, were examined as global indices of memory strength in recollection—one subjective and one objective—through two complementary analyses. Local voxel-wise univariate activity during recollection was correlated with each global index of memory strength, namely, objective whole-brain reactivation and subjective vividness ratings. These analyses yielded two maps of regions that covaried with reactivation and vividness of memories, respectively, and that were independent of the specific content (i.e. the specific video) comprising the recollection. A conjunction analysis revealed that the striatum, supplementary motor area, precentral gyrus, precuneus, superior parietal lobule, occipital pole, and right anterior hippocampus were positively correlated with memory indices whereas constituents of the default mode network yielded negative correlations (for a similar finding, see also, Cooper & Ritchey, 2019).

While informative, these analyses used a small sample and a relatively small set of stimuli and did not directly contrast vividness and reactivation maps. Here, we extended this work using a voxel-wise meta-analytic approach applied to data from three additional studies in addition to the study by St-Laurent and colleagues. Thus, we pooled over a much larger sample of participants (*N* =74) who performed similar cued retrieval tasks that used different stimuli. With this approach, we aimed to identify robust effects of vividness and whole-brain reactivation that generalize beyond any single study design. Additionally, beyond investigating similarity between the neural correlates supporting both measures of recollection, we contrasted the patterns revealed by each analysis to quantitatively test differences in neural correlates.

Computational models of episodic memory retrieval (Marr, 1971; McNaughton & Morris, 1987; Norman & O’Reilly, 2003) suggest that one major function of the hippocampus is pattern completion, a process that is a progenitor of cortical reinstatement of perceptual records that constitute a stored neocortical memory (Staresina & Wimber, 2019). Consistent with this idea, St-Laurent et al. (2015) found a within-subjects correlation between trial-to-trial univariate activity in the right anterior hippocampus and both whole-brain reinstatement and memory vividness. In the present meta-analytic study, we explored more fully how activity within this region relates to each measure. For each of the four studies, we correlated mean univariate hippocampal activity with local pattern classification evidence obtained via searchlight to determine if the hippocampus preferentially supported the reactivation of local cortical representations. In addition, to determine whether vivid memories are associated with cortico-hippocampal dynamics, we used a mediation analysis to assess whether univariate hippocampal activity promotes memory vividness indirectly by facilitating whole-brain reactivation.

Although vividness is a subjective measure and neural reactivation is an objective measure, both measures may be seen as indicating a kind of global “quality” of a memory, and it may be that reactivation is a prerequisite for vividly experienced memories (St-Laurent et al., 2015; for a demonstration using transcranial magnetic stimulation, see also, Waldhauser, Braun, & Hanslmayr, 2016). We therefore anticipated that voxels whose activity correlated with vividness judgements and neural reactivation would overlap spatially (as observed previously by St-Laurent et al., 2015), revealing areas in the brain participating in content-general recollective processes that supports global memory quality. Using a meta-analytic approach also allowed us to investigate the stability of these relationships across tasks and stimulus sets. Moreover, by pooling over a larger data set and directly contrasting correlates, we also expected to find differences that reflect the key distinction between subjective and objective measurement. First, because reactivation indexes the amount of perceptual information reinstated at retrieval, we expected to find associations in perceptual regions that typically process the representational content of an event. Next, because vividness judgement involves conscious monitoring and assessment of the reconstructed memory, we expected this measure to evoke the participation of higher order regions necessary for more executive processing. Finally, because the hippocampus is theorized to perform a pivotal role in recollection through pattern completion, we expected that hippocampal activation would relate to reactivation in cortical regions typically associated with representational content in memory, and that hippocampally-driven whole-brain reactivation would be associated with vivid remembering.

## 2. Method

### 2.1. Studies

The current meta-analysis was conducted using the data from four fMRI neuroimaging studies^1^ conducted by our group at the Rotman Research Institute at Baycrest Hospital. General study information is presented in Table 1. The demographic properties of the study samples were similar, with participants generally being well-educated young adults between the ages of 18 and 36.Acquisition parameters and behavioral analyses can be found in the original study publications, and a summary is presented in supplementary materials (Table S1).

**Table 1.**
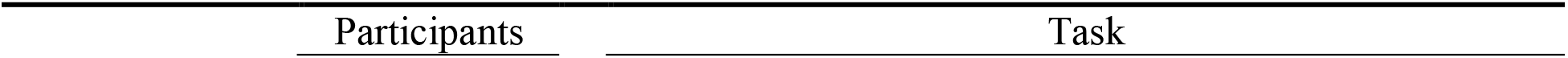
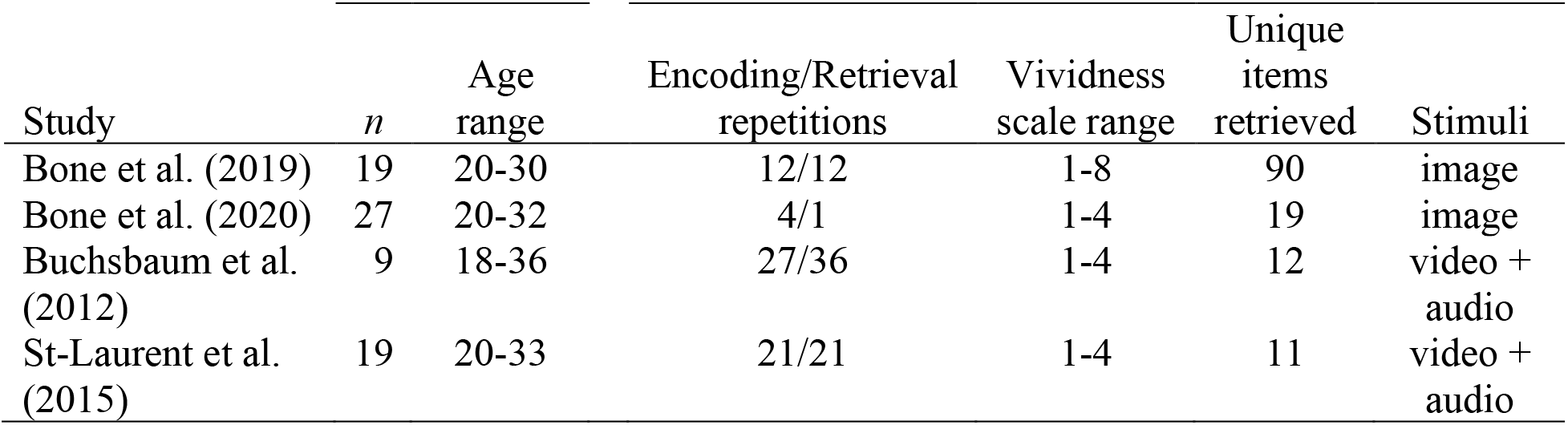
Design Information for Studies Included in Meta-analysis

| Study | *n* | Age range | Encoding/Retrieval repetitions | Vividness scale range | Unique items retrieved | Stimuli |
| --- | --- | --- | --- | --- | --- | --- |
| Bone et al. (2019) | 19 | 20-30 | 12/12 | 1-8 | 90 | image |
| Bone et al. (2020) | 27 | 20-32 | 4/1 | 1-4 | 19 | image |
| Buchsbaum et al. (2012) | 9 | 18-36 | 27/36 | 1-4 | 12 | video + audio |
| St-Laurent et al. (2015) | 19 | 20-33 | 21/21 | 1-4 | 11 | video + audio |

#### 2.1.1. Tasks

The studies included in the analysis varied in specific paradigms used, but all shared a general cued retrieval element. Across studies, the type of stimuli recalled also differed. Although all tasks employed visual stimuli, some included complex images whereas others used video clips with accompanying audio (see Table 1).

All the tasks contained an encoding phase where participants were presented with stimuli labeled with a visual word cue to encode and a retrieval phase where memory was tested. Multiple encoding and retrieval trials were performed on the same items to generate strong memories. Encoding was achieved by associating a word label or symbol with a relatively lengthy stimulus presentation (1.8 – 4.75 seconds range across picture studies; 3.6 – 9.0 seconds range across video studies). Retrieval was cued by briefly presenting the associated label, and reconstruction of the corresponding representation was performed during the presentation of a blank screen occupying a large time window (4.75 – 9.0 seconds range across studies). Following this recall, a screen asking participants to rate the vividness of their memory was presented (2.0 – 3.0 seconds across studies).

### 2.2. fMRI Preprocessing

Volumes were converted to NIFTI format and preprocessed using the Brain Imaging Data Structure (BIDS; Gorgolewski et al., 2016) fmriprep pipeline version 1.18 (Esteban et al., 2019; https://github.com/poldracklab/fmriprep). This workflow incorporates functionalities from standard fMRI processing software, including AFNI (v16.2.07), FSL (v5.0.9), ANTs (v2.1.0), and FreeSurfer (v6.0.1), to produce preprocessed volumes.

T1-weighted structural volumes were brain extracted using ANTs, tissue segmented with FSL’s FAST, and surface reconstructed with FreeSurfer. Volumes were then realigned to the Montreal Neurological Institute (MNI) template using a non-linear transformation implemented through ANTs.

Functional volumes were slice-timing corrected using AFNI’s 3dTshift to correct for temporal incongruency in volume slices. Each volume was then compared to a single reference volume to generate a rigid-body transform comprising six motion parameters (three translations and three rotations) using FSL’s MCFLIRT then applied using ANTs to correct for head motion. Lastly, a transformation was estimated using boundary-based registration in FreeSurfer to realign functional volumes, in native EPI space, to the T1-derived surface volumes. Final processed output in T1 space was used for univariate trial-wise beta estimation.

Mean global signal, mean tissue-specific signal, temporal, frame-wise displacement, rate of volume intensity change over time steps, and anatomical components via tCompCor and aCompCor were calculated and, with the six motion parameters previously estimated, used as nuisance regressors. The set of warps calculated to align T1 volumes to the MNI space was saved and applied to functional volumes for group level analyses.

To extract task related signal, a GLM was run for each subject over functional runs using a “least squares separate” beta estimation method (Mumford et al., 2012). The model included trial conditions for each event, convolved with the canonical hemodynamic response function, as well as a set of five nuisance regressors (constant, linear, quadratic, and higher order polynomial terms) to model low frequency noise components. The resulting trial-wise beta estimates were used for subsequent analyses.

#### 2.2.1. Multivariate Pattern Classification

A multivariate pattern classifier was trained on encoding trials to discriminate among stimuli using trial-wise beta images. The classifier was then used to “decode” neural activity from retrieval trials, that is, to identify the recalled stimuli, in order to assess the specificity of reactivation during recollection. First, a searchlight analysis was performed to determine participant-wise training features to be used for whole-brain classification. The searchlight used an 8mm radius and leave-one-out cross-validation to identify voxels that were most informative in predicting class labels for encoding data. Voxels in the upper 20% of area under the curve (AUC) values were selected to train the whole-brain pattern classifier that was trained on encoding trials and tested on retrieval trials.

Whole-brain classification models were generated using shrinkage discriminant analysis (SDA; Strimmer Lab, Imperial College London; strimmerlab.org/software/sda). This method is an extension of regularized linear discriminant analysis (Friedman, 1989) which addresses the issues inherent to linear discriminant classification when the number of predictors (voxels) is much larger than the number of observations (trials). These designs often violate the assumption of linear discriminant analysis that all categories’ covariance matrices are equal. James-Stein shrinkage reduces the variability in covariance matrices by shrinking values towards zero, ensuring that the inverse covariance matrix can be calculated for the discriminant function, and improves accuracy of classification.

SDA was performed for each participant and generated a discriminant score per stimulus for each retrieval trial used for classification. Classification yields posterior probabilities ranging from zero to one for every possible stimulus label, and a trial is assigned to the stimulus that obtains the highest probability. The posterior probability for the correct stimulus offers a meaningful and continuous measure of evidence of category membership per trial for each stimulus. The posterior probabilities (one per video) were used as strength of evidence measures for each class and a multi-class AUC score was computed using a one-against-all strategy. Specifically, the posterior probability for the “true” class was compared to the mean of the probabilities for all other classes, yielding a difference score for each memory retrieval trial. This process was repeated for each class and then averaged, yielding a final multi-class AUC.

### 2.3. First-level fMRI Analyses

#### 2.3.1. Local-global Analysis

A Local-Global Analysis (LGA) was performed to identify brain regions whose univariate activity was most strongly related to multivariate neural reactivation. LGA relates voxel-wise univariate activity (local) to whole brain pattern reactivation (global). This analysis was conducted via participant-wise multiple regression to regress trial-wise posterior probabilities corresponding to classification of the correct stimulus on to voxel-wise activity during retrieval. While St-Laurent and colleagues (2015) modeled the relationship with a linear model, the current analysis employed a quasibinomial logistic model, implemented in R (R Core Team, 2019) with the *glm* function, to better reflect the response distribution, which were the posterior probabilities of the classifier output for the “true” class, and were therefore ranged from 0-1. Variability in memory strength is known to be related to retrieval practice (e.g., Schuetze, Eglington, & Kang, 2019; Svoboda & Levine, 2009), and inter-stimulus variability may also be associated with fluctuations in the retrieved memories, therefore these influences were controlled for by including stimuli (video) and experiment block as nuisance covariates. This analysis was run using R formula syntax (i.e., reactivation ~ stimulus + block + voxel activation) where variables to the left of the ‘~’ represent the dependent variable and those to the right represent predictors. The output was whole brain maps for each participant that depicted where univariate brain activity covaried with the degree of reactivation of encoding activity at retrieval.

#### 2.3.2. Vividness Analysis

Univariate brain activity related to the vividness of retrieval was identified by regressing vividness ratings onto voxel-wise activity using multiple regression. As with the LGA, experiment block and stimulus were modeled as covariates of no interest. Since the 1-8 rating scale of Bone et al. (2019; see Table 1) was inconsistent with the 1-4 scale used in the other studies, ratings in this case were rebinned and then converted to a 1-4 scale. This relationship was modelled using ordinal logistic regression (rms package; Harrell, 2019) to maintain the ordered nature of the data whilst not assuming a metric scale across levels of vividness (R formula: vividness ~ stimulus + block + voxel activation).

#### 2.3.3. LGA-vividness Similarity

Memory vividness and neural reactivation are thought arise from general recollective processes, and therefore we anticipated that they should be associated with activity within overlapping sets of brain regions, as observed previously (St-Laurent et al., 2015). To identify this shared set of neural structures, the map-wise similarity between univariate patterns of activity associated with these two global measures of memory strength was computed via a whole-brain spatial correlation between the beta coefficient maps from the LGA and vividness analysis. The resulting coefficient represented the degree of spatial similarity between the two analyses.

To test whether distributed neural reactivation and trial-wise vividness judgments were related to one another, we correlated the two metrics via a quasibinomial regression model whereby reactivation probabilities were regressed onto vividness ratings while controlling for stimulus and experiment block (R formula: reactivation ~ stimulus + block + vividness).

#### 2.3.4. Hippocampal Modulation of Neural Reactivation and Memory Vividness

To reveal specific areas of the cerebral cortex where the level of reactivation covaried with the amplitude of hippocampal signal, local reactivation patterns identified via searchlight analysis (see section 2.2.1.) was regressed on to activity from a hippocampal region of interest (ROI) derived from the LGA using a quasibinomial model. Experimental block and stimulus were controlled for as covariates (R formula: searchlight reactivation ~ stimulus + block + hippocampus ROI activation).

The hippocampus is an important structure for memory retrieval, and reinstatement models predict a causal relationship whereby hippocampal activity precedes and supports cortical reactivation. A mediation model tested whether a relationship between vividness, reactivation, and the hippocampus was supported in our data by modelling vivid recollection as a function of hippocampally driven cortical reactivation. The effect of hippocampal activity on vividness was modelled indirectly through a mediation by whole-brain reactivation.

The mediation was modelled using a three-level mixed effect approach with subjects nested within studies (random intercept only) using the lme4 package in R (version 1.1.21; Bates, Maechler, Bolker, & Walker, 2015). A bootstrapping procedure was conducted to construct confidence intervals and assess the reliability of the indirect effect of hippocampus through reactivation on vividness (2000 bootstraps).

### 2.4. Meta-analysis

All meta-analyses were performed using a fixed-effects linear model implemented in the metafor package in R (version 2.1.0; Viechtbauer, 2010). A fixed-effects model assumes that study effects are sampled from a Gaussian distribution with some variance centered on a true population-level effect size. The model computes an estimate of the weighted effect across studies such that studies with a more reliable estimate of the true effect are weighted more strongly. Inverse weighting provides weights for each study by using the inverse of each effect’s sampling variance.

For the voxel-wise regression analyses (i.e., LGA, vividness, and hippocampus analysis), participant-level images were smoothed within individual brain masks (5 mm full width at half maximum Gaussian kernel). Study-level mean beta coefficients were used as a measure of effect and standard errors of the beta estimates were then calculated to be used for inverse weighting of the effects in the meta-analysis. Cochran’s *Q* statistic provides a measure of heterogeneity across studies and is computed from the summed squared deviation of each study effect from the weighted true effect estimate, while accounting for inverse weightings. The statistic is compared against a chi square distribution to assess reliability.

The relationship between trial-wise reinstatement evidence and vividness judgments was analyzed using mean study-level beta coefficients. Spatial similarity correlations from each study were transformed using Fisher’s *z* transformation and used as measures of effect size.

To determine reliable regions of contiguous voxel correlations, cluster-based thresholding was applied to correct for repeated comparisons. Spatial autocorrelation parameters were estimated from the residuals of a one-sample t-test for each study’s mean beta image. This provided an estimate of the inherent smoothness of the input images and was used to estimate an appropriate cluster-defining threshold using 3dClustsim in AFNI. Since we hypothesized that focal clusters would be found in the anterior hippocampus and whole-brain autocorrelation estimates are likely to yield cluster-defining thresholds that exceed the size of this structure, we used small volume correction to test for effects in this region. To achieve this, smoothness of the model residuals was estimated using a bilateral hippocampus mask based on ROIs from the Harvard Oxford atlas to determine a reasonable cluster threshold for hippocampal correlations.

Comparison of the LGA and vividness analysis was accomplished via linear contrast using a three-level mixed-effects model (random intercept only) with *lme4* in R. Participant-level images from both analyses were nested in participant and study to acknowledge the hierarchical structure of the data.

## 3. Results

### 3.1. Regions Associated with Vividness Judgements

Over all studies, vividness ratings of 1, 2, 3, and 4 were distributed across retrieval trials with proportions of 3.21% (*SD* = 1.79), 18.14% (*SD* = 3.13), 48.22% (*SD* = 2.83), and 30.42% (*SD* = 5.87), respectively (see Table 2). Most ratings were high (i.e., 3’s and 4’s) thus indicating that participants were successfully forming qualitatively strong and vivid mental representations of the cued stimuli.

**Table 2.** Average Proportion of Vividness Ratings by Study *Bone et al. (2019) used a rating scale of 1-8, therefore scores were binned and averaged to be comparable to the other studies.

| Study | Vividness rating |  |  |  |
| --- | --- | --- | --- | --- |
|  | 1 | 2 | 3 | 4 |
|  | *M (SD)* | *M (SD)* | *M (SD)* | *M (SD)* |
| Bone et al. (2019)* | 2.55 (4.02) | 18.73 (13.24) | 52.26 (11.78) | 26.46 (18.29) |
| Bone et al. (2020) | 2.93 (3.97) | 19.84 (12.20) | 47.98 (15.58) | 29.25 (21.71) |
| Buchsbaum et al. (2012) | 5.76 (2.65) | 20.43 (12.87) | 46.87 (18.60) | 26.94 (26.27) |
| St-Laurent et al. (2015) | 1.60 (1.66) | 13.57 (12.88) | 45.79 (18.45) | 39.03 (28.49) |

Meta-analysis of study-wise mean vividness beta images from the first-level vividness analysis, which capture the trial-wise relationship between vividness judgements and voxel activation, revealed a set of regions that were positively and negatively associated with vivid retrieval across studies (*p* <.05, *p* <.005 uncorrected, voxel-defining threshold = 50; Figure 1; Table 3). Positive associations between voxel activity and vividness judgements were found in the dorsal attention network. Regions included bilateral superior parietal lobules, intraparietal sulci, and frontal eye fields. Additionally, positive associations were found in frontal (i.e., left middle frontal gyrus and supplementary motor area), posterior (i.e., left posterior inferior temporal gyrus and ventral temporal cortices), and subcortical structures^2^ that spanned bilateral putamen and caudate nuclei. In contrast, negative associations with memory vividness were found mainly in constituents of the default-mode network. Activity in the ventromedial prefrontal cortex (vmPFC), cingulate, PCC, and angular gyri were all linked to reduced memory vividness. Other negative correlates included right dlPFC and mPFC, bilateral insular cortices, and right middle temporal gyrus.

**Figure 1.**
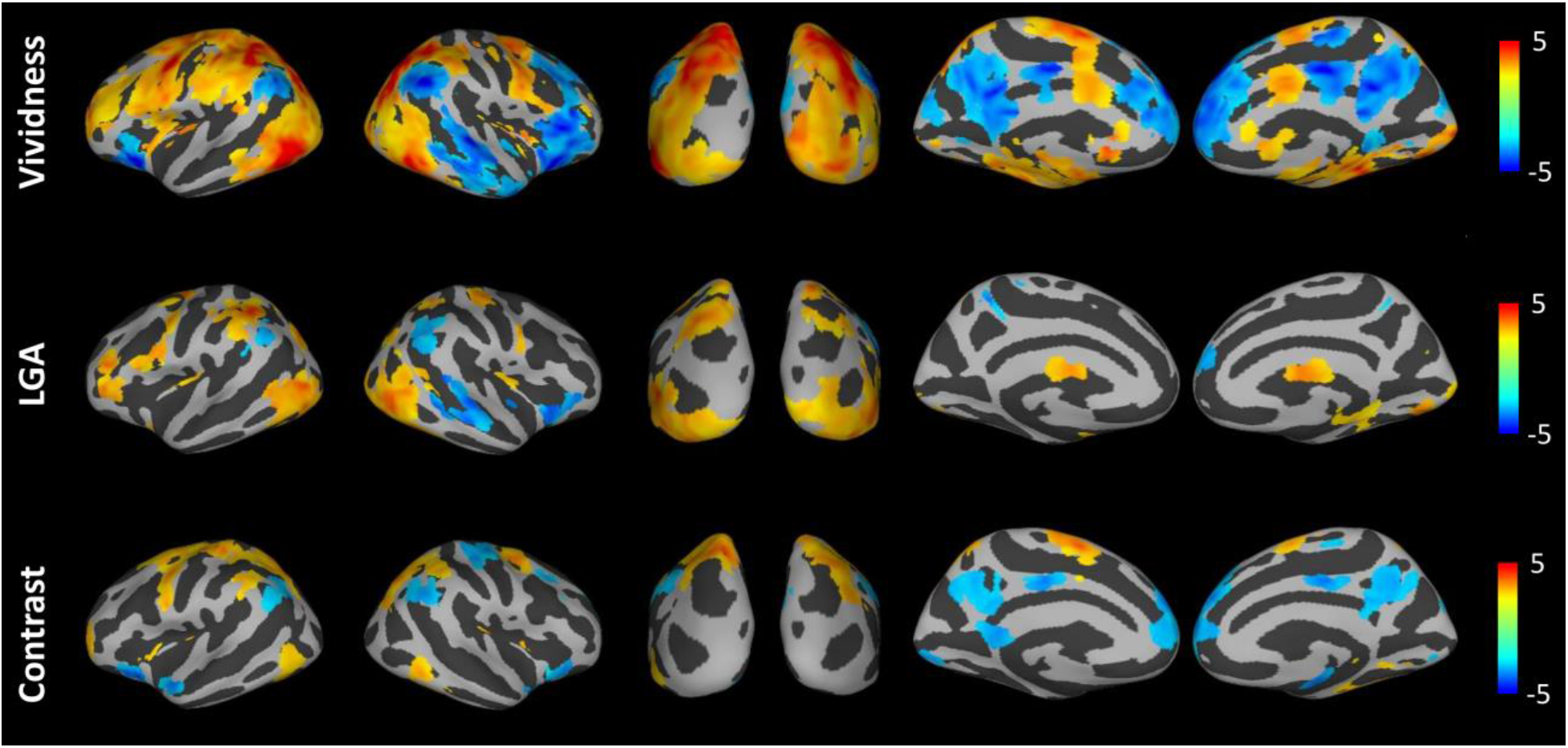
Regional Activity Predictive of Neural Reactivation and Vividness Judgements. For visualization, results are shown using a relaxed threshold (*p* <.01 uncorrected, cluster-defining threshold = 20). The LGA and vividness results (top and middle rows) depict *Z* scores and the contrast (bottom row; vividness - LGA) represents *t* statistics.

**Table 3.** Significant Clusters of Activity Found to Correlate with Vividness Judgements. Clusters are significant at p <.05, p <.005 uncorrected, using Monte Carlo simulation in AFNI’s 3dClustSim. MNI coordinates in millimetres. Volume indicate cluster size in voxels. ^a^ Hippocampus was examined with a small volume correction, significant at p < .05, p < .005 uncorrected, using Monte Carlo simulation in AFNI’s 3dClustSim.

| Region | Hemi. | Volume | Z value | MNI Coordinates |  |  |
| --- | --- | --- | --- | --- | --- | --- |
|  |  |  |  | x | y | z |
| Positive |  |  |  |  |  |  |
| Inferior temporal gyrus | R | 6612 | 9.12 | 53.80 | -61.80 | -10.20 |
| Intraparietal sulcus | R | 1203 | 8.67 | 30.40 | -75.80 | 41.70 |
| Putamen | L | 1176 | 8.50 | -23.50 | 3.70 | 9.80 |
| Putamen | R | 771 | 8.62 | 25.70 | 1.40 | 1.80 |
| Middle frontal gyrus | L | 318 | 5.69 | -42.20 | 59.90 | 5.80 |
| Precentral gyrus | R | 266 | 6.18 | 56.10 | -5.60 | 45.70 |
| Supplementary motor area | L | 249 | 6.46 | -2.40 | -10.30 | 61.70 |
| Precentral gyrus | R | 154 | 5.96 | 28.00 | -5.60 | 49.70 |
| Inferior occipital gyrus | L | 80 | 4.30 | -18.80 | -94.60 | -10.20 |
| Superior frontal gyrus | L | 55 | 5.04 | -23.50 | 55.20 | 5.80 |
| Hippocampus<sup>a</sup> | L | 17 | 4.30 | -16.40 | -10.30 | -18.20 |
| Hippocampus<sup>a</sup> | L | 8 | 4.30 | -25.80 | -8.00 | -26.10 |
| Hippocampus<sup>a</sup> | R | 8 | 3.51 | 16.30 | -10.30 | -18.20 |
| Negative |  |  |  |  |  |  |
| Medial prefrontal cortex | L | 1453 | -8.14 | -0.10 | 45.80 | 45.70 |
| Middle frontal gyrus | R | 706 | -8.15 | 44.40 | 24.80 | 41.70 |
| Angular gyrus | R | 500 | -7.51 | 60.80 | -59.50 | 25.70 |
| Posterior cingulate cortex | L | 498 | -6.57 | -2.40 | -47.80 | 25.70 |
| Middle temporal gyrus | R | 258 | -8.55 | 58.40 | -24.40 | -6.20 |
| Angular gyrus | L | 184 | -6.52 | -56.20 | -64.10 | 29.70 |
| Insula | L | 162 | -7.31 | -32.80 | 17.80 | -14.20 |
| Dorsal cerebellum | L | 129 | -5.23 | -28.10 | -85.20 | -38.10 |
| Cingulate cortex | R | 120 | -9.32 | 2.30 | -17.30 | 41.70 |
| Middle temporal gyrus | R | 91 | -5.52 | 46.70 | 13.10 | -42.10 |

Hippocampal correlations were determined using a small volume correction (*p* <.05, *p* <.005 uncorrected, voxel-defining threshold = 5). Significant positive clusters were found in bilateral anterior hippocampi (Figure 2), which is consistent with this region’s importance in vivid episodic memory retrieval.

**Figure 2.**
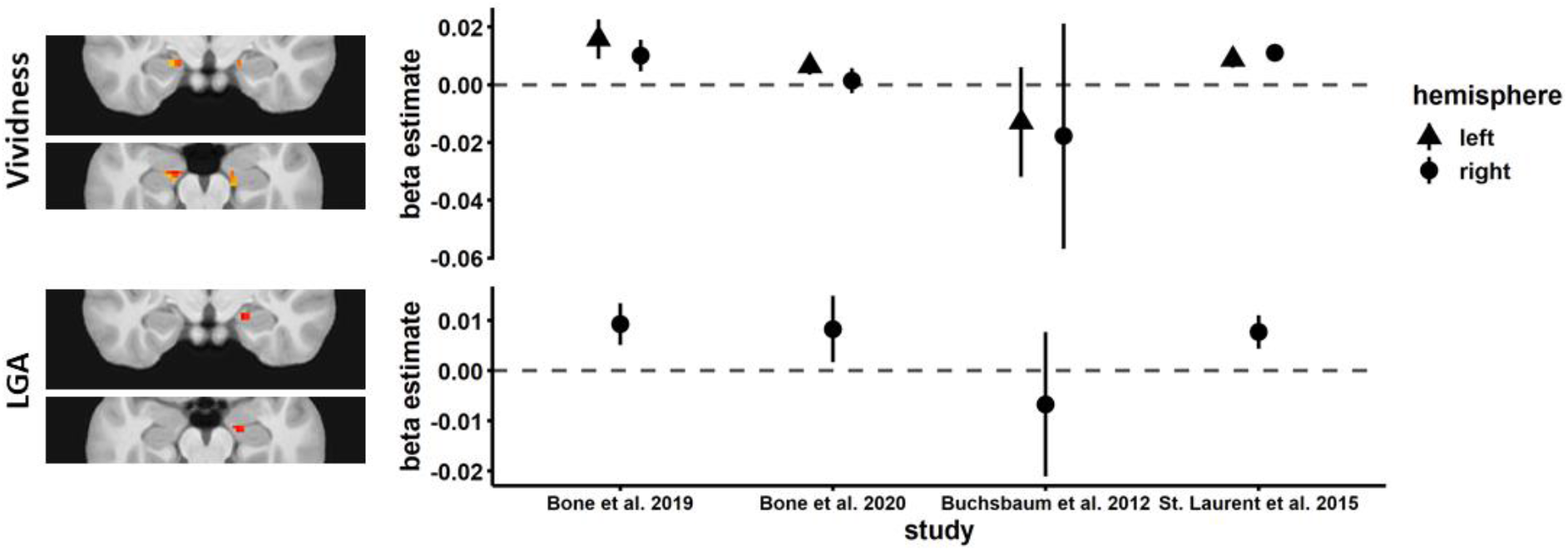
Hippocampus Predicting Neural Reactivation and Vividness Judgements. Significant clusters found to correlate with vividness judgements (top row; slice coordinates: y = −13, z = −20) and neural reactivation (bottom row; slice coordinates: y = −13, z = −19). Plots represent mean beta coefficients within each cluster by study. Error bars represent standard error.

### 3.2. Regions Revealed by LGA to be Associated with Whole-brain Reactivation

Whole-brain classifiers trained on perceptual activity at encoding and tested on retrieval activity performed above chance levels for all studies (one-sample tests, *p* <.05), indicating successful stimulus-specific decoding (see Table 4).

**Table 4.** Classifier Performance by Study. Performance was quantified with area under the curve values with chance (.5) centered on zero. *Denotes a significant difference from chance.

| Study | $M (SD)$ | $t$ value | |
| --- | --- | --- | --- |
| Bone et al. (2019) | 0.05 (0.03) | 6.11 | * |
| Bone et al. (2020) | 0.01 (0.01) | 2.06 | * |
| Buchsbaum et al. (2012) | 0.06 (0.07) | 2.92 | * |
| St-Laurent et al. (2015) | 0.12 (0.05) | 9.84 | * |

The LGA identified sets of regions whose activation was positively associated with whole-brain reactivation across studies and stimuli (*p* <.05, *p* <.005 uncorrected, voxel-defining threshold = 50; Figure 1; Table 5). Notably, a strong relationship with neural reactivation was detected within the early visual cortex. Regions in the dorsal attention network also correlated with reactivation (i.e., left intraparietal sulci, and frontal eye field), in addition to temporal sites (i.e., right middle temporal gyrus, right ventral temporal cortex) and left ventral putamen. A negative relationship with reactivation was only evident in the right middle temporal gyrus.

**Table 5.**
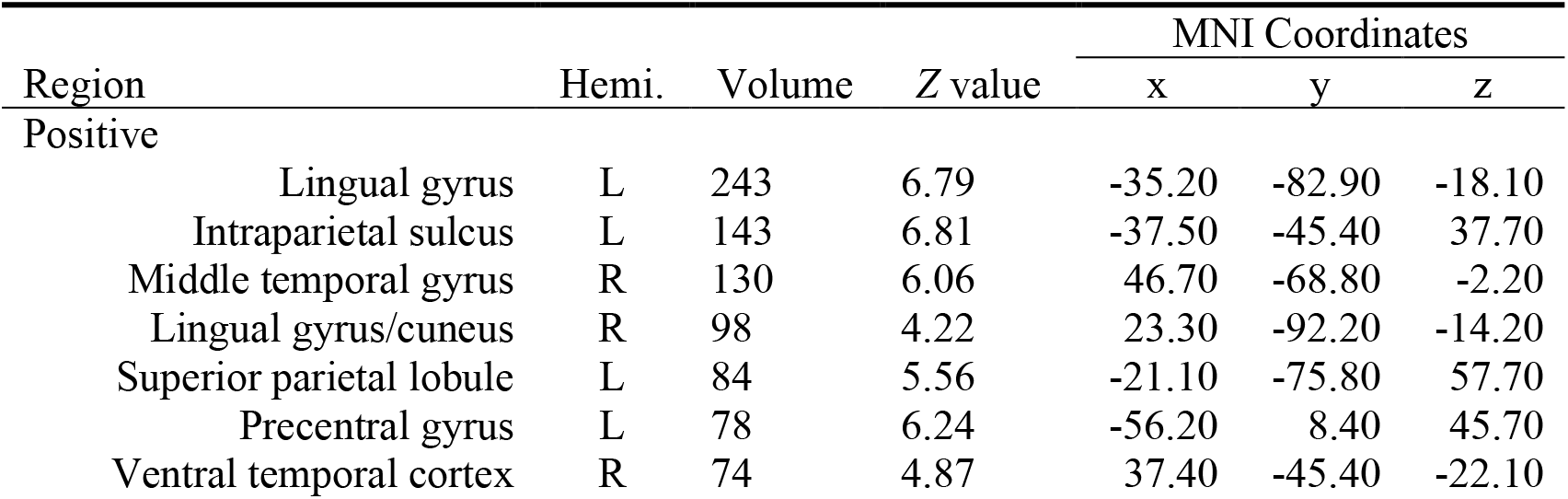
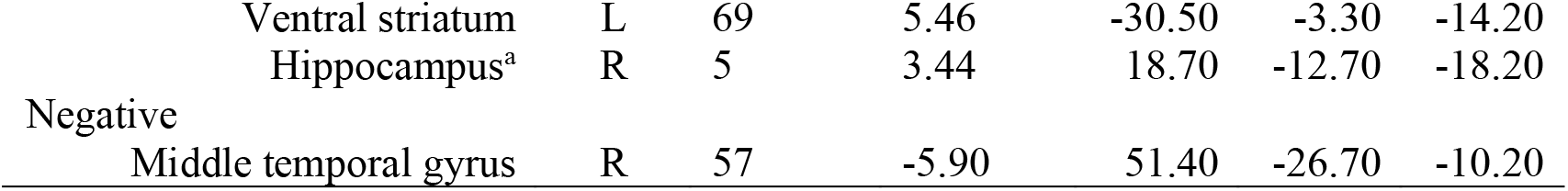
Significant Clusters of Activity Found to Correlate with Neural Reactivation. Clusters are significant at p < .05, p < .005 uncorrected, using Monte Carlo simulation in AFNI’s 3dClustSim. MNI coordinates in millimetres. Volume indicate cluster size in voxels. ^a^Hippocampus was examined with a small volume correction, significant at p < .05, p < .005 uncorrected, using Monte Carlo simulation in AFNI’s 3dClustSim.

| Region | Hemi. | Volume | Z value | MNI Coordinates |  |  |
| --- | --- | --- | --- | --- | --- | --- |
|  |  |  |  | x | y | z |
| Positive |  |  |  |  |  |  |
| Lingual gyrus | L | 243 | 6.79 | -35.20 | -82.90 | -18.10 |
| Intraparietal sulcus | L | 143 | 6.81 | -37.50 | -45.40 | 37.70 |
| Middle temporal gyrus | R | 130 | 6.06 | 46.70 | -68.80 | -2.20 |
| Lingual gyrus/cuneus | R | 98 | 4.22 | 23.30 | -92.20 | -14.20 |
| Superior parietal lobule | L | 84 | 5.56 | -21.10 | -75.80 | 57.70 |
| Precentral gyrus | L | 78 | 6.24 | -56.20 | 8.40 | 45.70 |
| Ventral temporal cortex | R | 74 | 4.87 | 37.40 | -45.40 | -22.10 |

**Table 5.** Significant Clusters of Activity Found to Correlate with Neural Reactivation. Clusters are significant at $p < .05$ , $p < .005$ uncorrected, using Monte Carlo simulation in AFNI's 3dClustSim. MNI coordinates in millimetres. Volume indicate cluster size in voxels.
|  |  |  |  |  |  |  |  |
| --- | --- | --- | --- | --- | --- | --- | --- |
|  | Ventral striatum | L | 69 | 5.46 | -30.50 | -3.30 | -14.20 |
|  | Hippocampus<sup>a</sup> | R | 5 | 3.44 | 18.70 | -12.70 | -18.20 |
| Negative | Middle temporal gyrus | R | 57 | -5.90 | 51.40 | -26.70 | -10.20 |

Consistent with the vividness analysis, right anterior hippocampal activity reliably predicted greater whole-brain reactivation (Figure 2; *p* <.05, *p* <.005 uncorrected, voxel-defining threshold = 5).

### 3.3. Shared Neural Correlates between Reactivation and Vividness

Consistent with the hypothesis that neural reactivation is necessary for vividly experienced memories, a positive relationship was found between trial-wise memory vividness and whole-brain neural reactivation across studies, *Z* =3.86, *p* < .001, *Q*(3) = 19.55, *p* <.001.

There is evidence that neural reactivation and memory vividness, as summary measures of global memory content in recollection, rely on overlapping neural correlates (Bone et al., 2019; Dijkstra et al., 2017; St-Laurent et al., 2015; Wing et al., 2015). To examine to what extent the measures were associated with similar sets of brain regions, spatial congruence was assessed between the two analyses. Pearson correlations between beta coefficient maps from the LGA and vividness analysis were reliable across studies, *r* =.14, *Z* =6.42, *p* < .001, *Q*(3) = 9.97, *p* = .02, indicating that neural reactivation and vividness ratings were predicted, in part, by shared neural activity.

A conjunction analysis (Figure 3) visualized areas of overlap between analyses. This conjunction revealed that dorsal attention regions in the superior parietal lobe were consistently related to perceived vividness and brain reactivation at retrieval. Correlations with the left posterior inferior temporal gyrus, right posterior ventral temporal cortex, and left putamen were also found to overlap between measures. Notably, these regions were all found to relate positively to both measures, thus representing a common set of regions that are supportive of successful memory retrieval in the studies considered.

**Figure 3.**
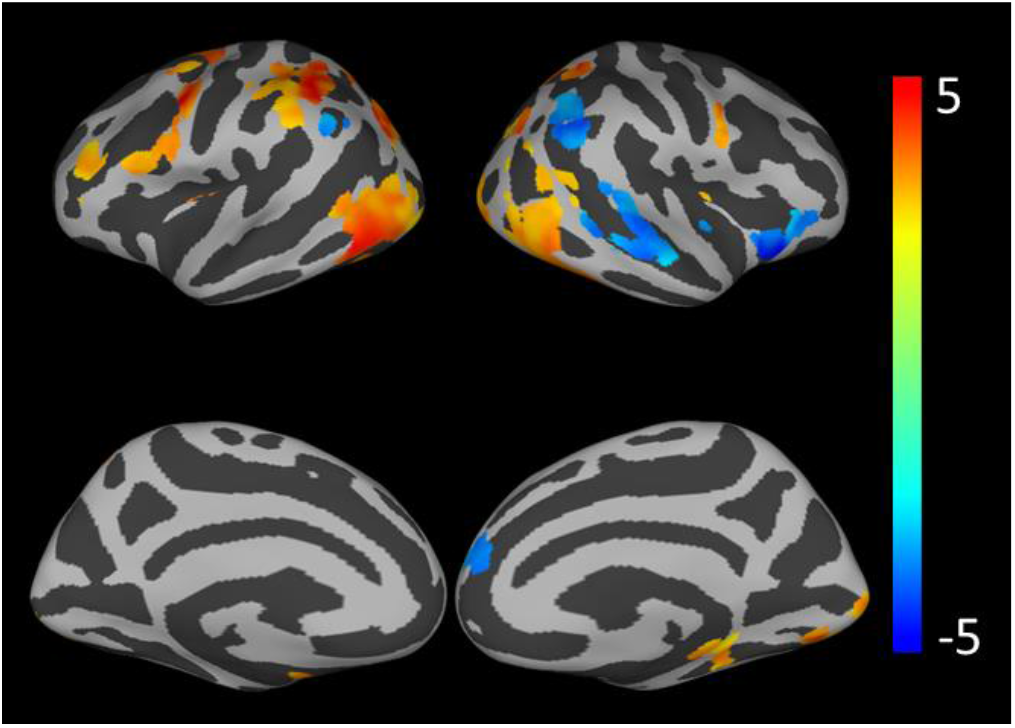
Conjunction of LGA and Vividness Analysis. Cluster depict *Z* scores. For visualization, the conjunction was computed using a relaxed threshold (*p* < .01 uncorrected, cluster-defining threshold = 20) for the LGA and vividness analysis. *Z* scores within the conjunction were calculated as the average between the LGA and vividness analysis results.

### 3.4. Comparing Vividness-related and LGA Responses

A linear mixed-effects model was used to estimate statistical contrasts between neural activity associated with vividness ratings and neural reactivation (Figure 1; Table 6). Greater correlations with vividness ratings were primarily observed in the dorsal attention network and the striatum, bilaterally. Furthermore, there was less of a negative relationship with neural reactivation throughout the default mode network (i.e., posterior cingulate cortex, angular gyrus), whereas vividness ratings were more reliably modulated by suppressed activity in these regions. Reactivation, however, was found to have a stronger positive relationship with right hippocampal activations. Additionally, stronger correlations with neural reactivation were observed in early visual regions, though this finding did not survive cluster correction.

**Table 6.**
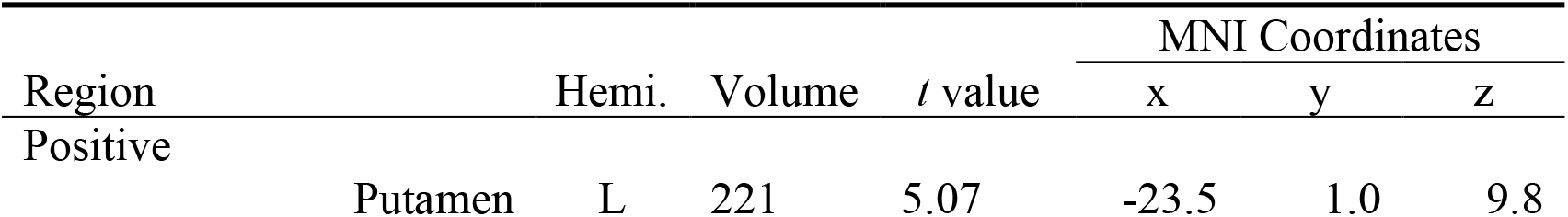
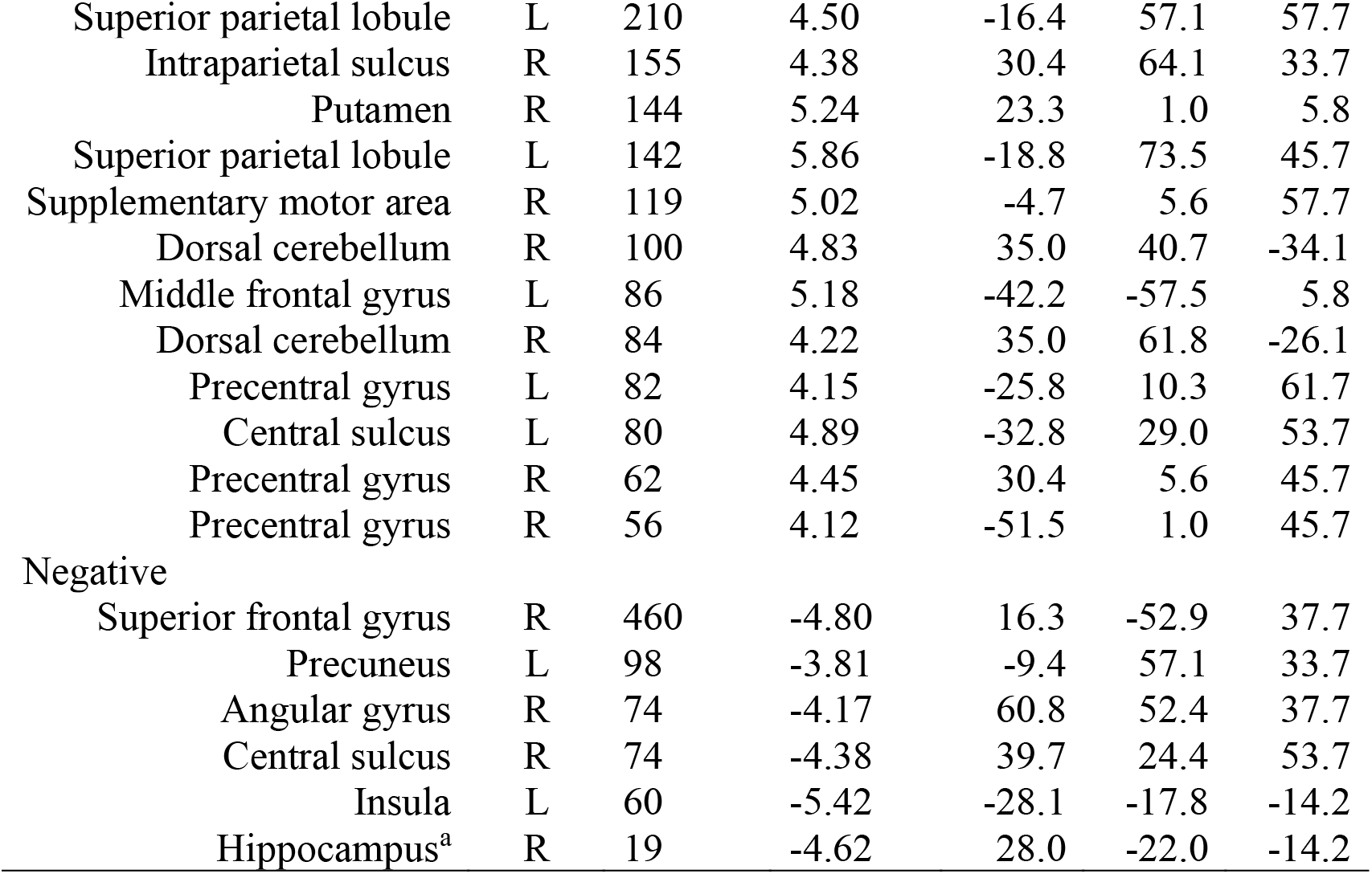
Regions that Significantly Differed between the LGA and Vividness Analysis. Clusters are significant at p < .05, p < .005 uncorrected, using Monte Carlo simulation in AFNI’s 3dClustSim. MNI coordinates in millimetres. Volume indicate cluster size in voxels. ^a^ Hippocampus was examined with a small volume correction, significant at p < .05, p < .005 uncorrected, using Monte Carlo simulation in AFNI’s 3dClustSim.

In addition to regional differences in correlation strength between activation and each measure, there was a difference in how widespread and distributed the results were. Correlates of vividness ratings were more spatially extensive than those identified by the LGA. Interstudy differences may have introduced different amounts of variability among correlates between the two measures, and this was explored via pairwise spatial correlations for each result (i.e., vividness analysis and LGA) between each study mean beta image.

The correlation matrix is presented in Figure 4 and reveals a pattern of weaker correlations between studies for LGA results and stronger, positive correlations for vividness analyses. Interestingly, despite weaker correlations for LGA images, the strongest correspondence was found between the two studies (Buchsbaum et al., 2012, St-Laurent et al., 2015) that used multimodal video stimuli. The pattern of correlations for each measure was tested using Steiger’s chi square test (Steiger, 1980), and found to reliably differ, *Χ*^2^(6) = 25276.96, *p* < .001. This suggests that the neural correlates associated with vividness ratings were more consistent across studies, but regions relevant for neural reactivation were influenced to a greater extent by differences in design and or stimuli.

**Figure 4.**
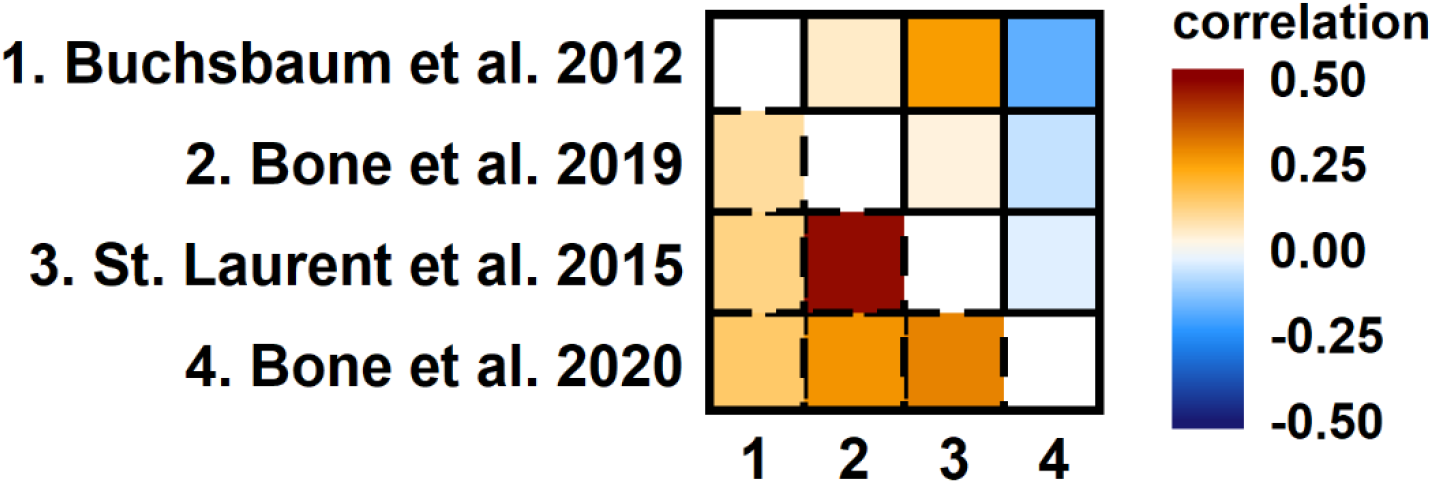
Study-wise Correlations of LGA and Vividness Analysis Results. Correlations between vividness maps appear in the lower triangle (dashed border) and correlations between reactivation maps appear in the upper triangle (solid border).

### 3.5. Hippocampally-associated Reactivation

The relationship between hippocampal activation and local patterns of neural reactivation at retrieval was explored by correlating mean activity in the hippocampal cluster identified in the LGA with searchlight-based reactivation evidence (Figure 5; see Table 7 for cluster report). Hippocampal activity predicted greater local reactivation in regions linked to informational representation in the visual (i.e., ventral temporal cortex, parahippocampal gyrus) or cross-modal (i.e., precuneus, angular gyrus) domains. Reduced hippocampal activation, however, was associated with greater reactivation in the mPFC.

**Figure 5.**
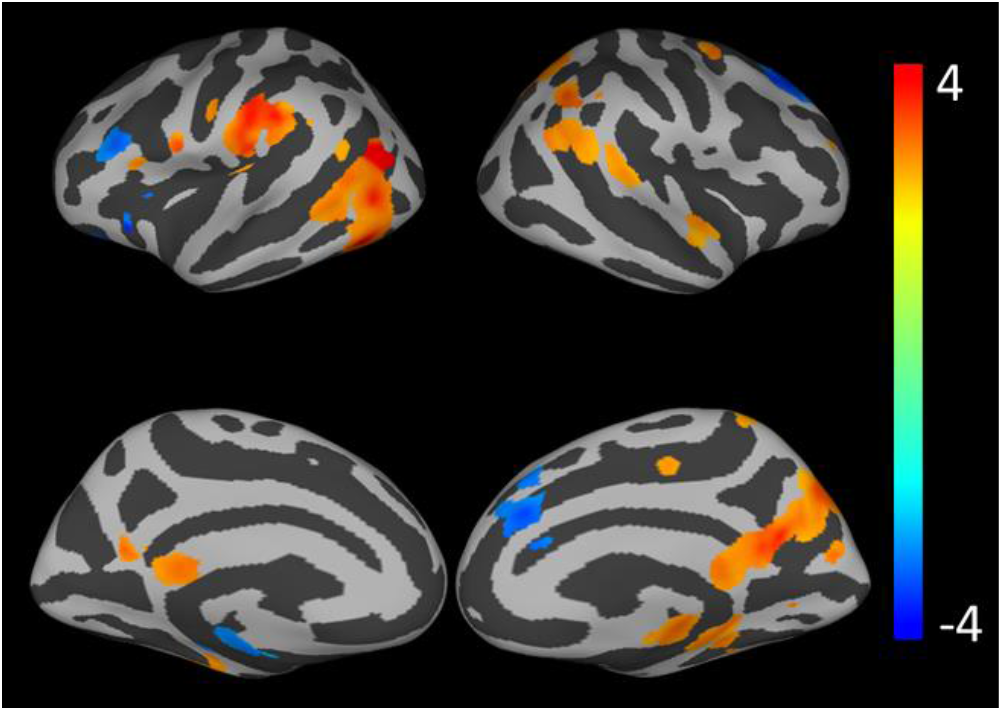
Regional Reactivation Predicted by Right Hippocampus Activity. Values depict *Z* scores. For visualization, results are shown using a relaxed threshold (*p* < .01 uncorrected, cluster-defining threshold = 20).

**Table 7.** Regional Reactivation Significantly Predicted by Right Hippocampus. Clusters are significant at p < .05, p < .005 uncorrected, using Monte Carlo simulation in AFNI’s 3dClustSim. MNI coordinates in millimetres. Volume indicate cluster size in voxels. *A cluster in the posterior parahippocampal gyrus (48 voxels) approached the voxel-defining threshold of 50 voxels. It is likely that such a large threshold is too stringent to detect relationships in a circumscribed region such as the posterior parahippocampal gyrus, therefore this cluster is included in the table.

|  |  |  |  | MNI Coordinates |  |  |  |
| --- | --- | --- | --- | --- | --- | --- | --- |
| Region |  | Hemi. | Volume | Z value | x | y | z |
| Positive |  |  |  |  |  |  |  |
|  | Superior parietal lobule/precuneus | R | 300 | 5.65 | 4.60 | -64.10 | 17.80 |
|  | Middle temporal gyrus | L | 189 | 5.24 | -39.80 | -57.10 | -2.20 |
|  | Supramarginal gyrus | L | 143 | 4.36 | -67.90 | -31.40 | 25.70 |
|  | Inferior parietal lobule | R | 90 | 4.33 | 53.80 | -54.80 | 45.70 |
|  | Parahippocampal gyrus* | R | 48 | 3.60 | 23.30 | -33.70 | -10.20 |
| Negative |  |  |  |  |  |  |  |
|  | Dorsomedial prefrontal cortex | R | 163 | -4.64 | 14.00 | 24.80 | 41.70 |

The theorized role of the hippocampus in performing cortical pattern completion in support of successful recollection was tested using a mediation model. The hippocampus is implicated in cortical reactivation during retrieval, therefore we supposed that the relationship between neural reactivation and experienced vividness is such that hippocampally-driven reactivation indirectly influences vividness judgements. This hypothesis was supported by a significant indirect effect, *β* =.007, *SE* =0.13, 95% CI [0.002, 0.01], however a reliable direct effect of hippocampus activity on vividness judgments was also observed, *β* =.02, *SE* =0.29, 95% CI [0.008, 0.03], indicating a partial mediation. This result suggests that the hippocampal results observed in the vividness analysis and LGA reflect this region’s role in modulating reactivation throughout the brain, which enables richly perceived mnemonic representation.

## 4. Discussion

The present study investigated systematic relationships between global indices of memory quality and their associated patterns of neural activation. We predicted that whole-brain reactivation and memory vividness, because they are both known indicia of a recollective process, should involve similar underlying neural circuitry. This prediction was confirmed with a conjunction analysis, and by positive spatial correlations between the LGA and vividness analysis, replicating the findings originally reported by St-Laurent and colleagues (2015). The existing literature has established a clear positive relationship between encoding-trained, retrieval-tested classifier evidence and memory vividness (e.g., Bone et al., 2019; Dijkstra et al., 2017; Wing et al., 2015) and the current result suggests that these measures are supported by some of the same brain networks. However, we also identified important differences in the neural correlates of reactivation and memory vividness that offer insight into how objective and subjective measures of recollection may diverge. Differences have previously been shown between objective and subjective measures of memory (e.g., Richter et al., 2016; Thakral, Madore, & Schacter, 2020) but, to our knowledge, this is the first attempt to directly contrast reactivation and vividness judgements.

Positive associations were found between dorsal attention network activation and both measures of memory quality. The superior parietal lobule has been well-studied for its role in top-down attention (Corbetta & Schulman, 2002), and has been suggested to perform a similar top-down role in episodic retrieval (Guerin, Robbins, Gilmore, & Schacter, 2012). Furthermore, previous literature has linked the precuneus, a subregion of the superior parietal cortex, to vivid remembering (Fletcher et al., 1995; Fulford et al., 2018; Gilboa, Winocur, Grady, Henevor, & Moscovitch, 2004; Lundstrom, Ingvar, & Petersson, 2005; Sreekumar, Nielson, Smith, Dennis, & Sederberg, 2018), a phenomenon that may be linked to egocentric processing of visual perspective (St Jacques, Szpunar, & Schacter, 2017). Vivid recollection relies on top-down processes to maintain and construct rich mnemonic representations, and the studies we analyzed employed paradigms in which participants were encouraged to construct vivid representations of complex visual stimuli.

The frontal eye fields were also associated with greater memory as indicated by vividness judgements and neural reactivation. This structure has historically been discussed in terms of its role in oculomotor control, however, emerging evidence suggests frontal eye field involvement in memory processes. Indirect anatomical pathways have been discovered in macaques from the medial temporal lobes to oculomotor regions (Shen, Bezgin, Selvam, McIntosh, & Ryan, 2016), and a functional pathway has similarly been identified with human simulations (Ryan et al., 2018; Spiegler, et al., 2016). This pathway is thought to allow past experiences to influence one’s visual exploration of the environment, an interpretation supported by evidence that the reinstatement of eye movements deployed at encoding facilitates successful retrieval (Bone et al., 2019; Wynn, Shen, & Ryan, 2019). The studies included in the current meta-analysis used visually complex stimuli that should promote gaze fixation reinstatement at retrieval. A positive association with memory vividness and neural reactivation in the frontal eye fields may reflect spatial representations encoded at perception guiding oculomotor behavior when reconstructing a visual representation. Therefore, the association between dorsal attention regions and measures of global memory quality may indicate that attention applied to retrieve perceptual details could enhance the fidelity of the mnemonic reconstruction.

Importantly, regional differences were observed when the LGA and vividness analyses were contrasted. The clearest distinction between the two results maps was a tendency for memory vividness to be more strongly related to increased activity in the dorsal attention and suppressed activity in the default mode networks. This difference highlights the subjective and conscious nature of vividness as a measure of memory quality. Reporting the perceived detail of a mental representation requires an effortful application of attention and should be expected to tax controlled retrieval processes. For example, working memory processing is necessary to maintain and manipulate representations as a visual mental image is constructed (Baddeley & Andrade, 2000; Engelhard, van den Hout, & Smeets, 2011; Keogh & Pearson, 2014), and the superior parietal lobule is known to support these operations (e.g., Koenigs, Barbey, Postle, & Grafman, 2009; Nee et al., 2013). Alternatively, evidence suggests that particularly strong memories may capture bottom-up attention, and activity associated with vivid retrieval in the posterior parietal cortex, particularly regions extending ventrally, may reflect this (Cabezza, Ciaramelli, & Moscovitch, 2012; Sestieri, Shulman, & Corbetta, 2017). Regardless, increased activation in posterior parietal regions when memories were most vivid is indicative of recruitment of visual attentional processes (see, Ciaramelli, Grady, & Moscovitch, 2008), and a greater engagement of these processes may be a critical aspect that dissociates memory vividness from neural reactivation as a subjective, rather than objective, measure of global memory quality.

The striatum was positively correlated with both vividness judgments and neural reactivation at retrieval. This finding has been interpreted as reflecting sequential processing when video stimuli were used (St-Laurent et al., 2015), however the current meta-analysis suggests that this relationship holds even for pictorial stimuli that lack explicit sequence information. It is possible that in such cases sequential information may still be represented as visual representations are thought to be reconstructed through a sequence of attentional foci, but a stronger relationship with vividness judgements than reactivation indicates such a general interpretation cannot fully account for striatal involvement.

The structures comprising the striatum have been noted for their response to reward (Balleine, Delgado, & Hikosaka, 2007; Haber 2011). A hypothesis for striatal involvement during retrieval suggests these regions track reward value, just as they have been found to do in other domains (Han, Huettal, Raposo, Adcock & Dobbins, 2010). From this perspective, the striatum is key in maintaining task goals and exhibits responsiveness when outcomes are rewarding or align with the perceived objective. In the context of recognition tasks, successful retrieval is implicitly rewarding, even when no concrete reward or reinforcement is present. This interpretation is consistent with studies that report a pattern of greater striatal activity for hits (Han et al, 2010; Spaniol et al., 2009; von Zerssen, Mecklinger, Opitz, & von Cramon, 2001) and for false alarms (Abe et al., 2008; von Zerssen et al., 2001) than correct rejections, indicating that subjectively perceived retrieval success engages these structures even during objective retrieval failures. Moreover, the same pattern has been shown to amplify when a monetary value was assigned to hit trials and to reverse when correct rejections were incentivized (Han et al., 2010). From this perspective, striatal activity at retrieval tracks subjective perception of task success and is closely linked to goal-directed behavior. Memory vividness judgements provide a subjective index of memory retrieval success during recall. Although previous work has only implicated caudate activity in goal attainment, while the contrast between reactivation and vividness judgements revealed differences in clusters within the putamen that extend into caudate, stronger striatal activation when vivid memories are retrieved may reflect a reward response elicited by meeting task goals.

Some regions were found to be more readily associated with neural reactivation than with vividness. A cluster in the central sulcus, and a strong effect in left visual cortex that was too localized to survive cluster correction (local peak *t*-value = −4.01, cluster size = 27 voxels), both suggest that whole-brain reactivation was more closely related to activity in early sensory cortices. Perception of complex stimuli engenders processing of their constituent features, starting in the earliest sensory cortices and feeding upwards to generate ever more complex representations. It is perhaps low-level features, representing specific perceptual details, that may best discriminate between complex stimuli that share higher-level, semantic features. Thus, stronger neural reactivation will naturally be found when structures early in the processing stream are engaged. Indeed, early visual perceptual activity has been reported to be important in classifying the identity of retrieved representations (Bone et al., 2019, 2020; Bosch, Jehee, Fernández, & Doeller, 2014; Buchsbaum et al., 2012; Lohnas et al., 2017; St-Laurent et al., 2015) and activity in these regions has been linked to true recognition (Slotnick & Schacter, 2004). Activity in early sensory structures may even be elicited in the absence of conscious awareness (Frey et al., 2016; Slotnick & Schacter, 2004, 2006), a phenomenon that may dissociate the mechanisms underlying neural reactivation, as an objective measure of recollection, from those relevant to subjective measures like vividness judgements.

The neural correlates of reactivation and vividness also differed in their inter-study variability. Inter-study correlations indicated that the regions associated with vividness judgments were related positively across studies. Reactivation correlates, however, were less consistent between studies. This pattern suggests that a general set of brain structures is associated with vivid memories, whereas many regions that are related to reactivation are specific to the parameters and stimulus sets used by a particular study. That is, the properties of stimuli are likely an important factor in determining which structures drive neural reactivation. This observation might explain why the neural correlates of reactivation that emerged from the meta-analysis were sparser than those of vividness ratings, and did not include negative associates throughout the default mode network. Negative correlations in the default mode network were a prominent finding in the original analysis by St-Laurent and colleagues (2015), and this finding was interpreted as reflecting mind wandering interfering with mental replay. Perhaps, the relationship between default mode network activity and memory reactivation differs as a function of task parameters such as stimulus complexity and the amount of exposure and retrieval opportunities. As St-Laurent et al. (2015) found a negative relationship between default mode network activity and whole-brain reactivation using video stimuli, mind wandering may affect representational specificity to a greater extent when mental representations are more complex and highly dimensional. Vividness judgements, however, appear to be reliably affected by mind wandering despite task design differences. Presumably, the interference with top-down attention that occurs with mind wandering competes for attentional resources that promote vivid recollection.

We hypothesized that hippocampal activity would reliably predict both our measures of global memory quality. The hippocampus is a key structure involved in episodic memory and is theorized to interact with the cortex during retrieval through pattern completion and subsequent neocortical reactivation (Marr, 1971; McNaughton & Morris, 1987; Norman & O’Reilly, 2003). Thus, we expected that hippocampal activity would covary with the emergence of activity patterns that contain representational information specific to the retrieved stimulus. Previous work has reported correlations between hippocampal activity and reactivation in localized cortical ROIs (Bosch et al., 2014; Ritchey et al., 2013; Staresina et al., 2012; Tompary, Duncan, & Davachi, 2016), but the LGA meta-analysis extended these findings to include whole-brain reactivation patterns. The vividness meta-analysis also indicated that pattern completion processes performed by the hippocampus are related to the phenomenology of remembering that accompanies recollection.

When we correlated univariate anterior hippocampal activity with reactivation across the cortex, we observed stronger correlations in posterior cortical sites. The hippocampus is thought to perform a critical role in memory via pattern completion processing (Marr, 1971; McClelland, McNaughton, & O’Reilly, 1995), and it is understood to trigger reactivation within the sensory processing hierarchy (for a review see, Staresina & Wimber, 2019). Consistent with this interpretation, hippocampal activity has been associated with reinstatement in the visual system when visual stimuli are retrieved (e.g., Bosch et al., 2014; Mack & Preston, 2016). Similarly, we found that hippocampal activity was related to reinstatement in regions known to process representational content of visual stimuli (i.e., parahippocampal gyrus and ventral temporal cortex). We also detected hippocampally-linked reactivation in the angular gyrus, superior temporal gyrus, and precuneus which have been linked to higher-order perceptual processing. Multimodal integration has been proposed to occur in the angular gyrus (Shimamura, 2011; Tibon et al., 2019) and superior temporal gyrus (Beauchamp, Lee, Argall, & Martin, 2004; Metzger et al., 2020) while the precuneus has been linked to similar integrative processing (Naghavi & Nyberg, 2005) in addition to spatial, self-referential processes that are key for episodic memory representation (Cavanna, 2007; Hebscher, Ibrahim, & Gilboa, 2020). A recent effective connectivity analysis found that anterior hippocampus-precuneus connectivity modulated subjective memory confidence, suggesting that it may support automatic, unconscious processes that contribute to metacognitive memory appraisal (Ren et al., 2018). Neural reactivation is likely one such automatic process that facilitates conscious memory recollection.

Our mediation analysis also revealed that reactivation partially mediated the effect of anterior hippocampus activity on vividness judgements. The indirect effect supports the hypothesis that the hippocampus performs pattern completion to reinstate cortical activity at retrieval, which enables one to subjectively re-experience a memory in vivid details. The same relationship has been demonstrated previously with graded old/new judgements as a behavioral measure of recognition memory success (Ritchey, Wing, LaBar, & Cabeza, 2013). Our results provide further evidence that hippocampally-driven reactivation influences conscious recollective processes engaged during vividness judgements.

## 5. Conclusions

The current investigation offers novel evidence that neural reactivation patterns and memory vividness ratings are partly supported by separable neural mechanisms. Despite both measures demonstrating correlated patterns of distributed coactivation, vividness judgements were more strongly associated with regions implicated in attentional processing and perceived task success. Reinstatement, however, was predicted more strongly by early sensory activity. As a subjective measure of memory, vividness judgements are fundamentally tied to one’s conscious experience. By contrast, neural reactivation is a purely objective measure that involves a “third-party” readout of brain activity patterns. Thus, while the extent of neural reinstatement may be consciously conveyed by vividness reports, the neural processes that give rise to reinstatement need not always be consciously accessible, and the dissociation between the neural correlates of these measures reflects this key distinction.

## Acknowledgements

This work was supported by an NSERC discovery award (RGPIN 386631-10) and CIHR project award (PJT152879) to Bradley R. Buchsbaum

## Credit Authorship Statement

**Ryan M. Barker:** Formal analysis, Methodology, Writing – original draft, Writing – review & editing, Visualization. **Marie St-Laurent:** Methodology, Writing – review & editing. **Bradley R. Buchsbaum:** Conceptualization, Methodology, Writing – review & editing, Supervision, Project administration, Software, Resources, Funding acquisition.

## Data Availability

All data is available on request via email directed to Bradley R. Buchsbaum.

## Supplementary Materials

**Table S1.** Summary of Imaging Parameters Used by Studies Analyzed

|  | Head<br>coil | FOV (cm) | Matrix<br>size | In-plane<br>resolution<br>(mm) | Slice<br>Thickness<br>(mm) | Flip<br>angle | TE<br>(ms) | TR |
| --- | --- | --- | --- | --- | --- | --- | --- | --- |
| Bone et al.<br>(2019) | 12-channel | 22.5 x 22.5 | 96 x 96 | 2.35 x 2.35 | 3.5 | 62° | 27 | 1.5 s |
| Bone et al.<br>(2020) | 32-channel | 22 x 22 | 110 x 110 | 2.0 x 2.0 | 2.0 | 62° | 30 | 1.77 |
| Buchsbaum<br>et al.<br>(2012) | 32-channel | 22.5 x 22.5 | 96 x 96 | 2.5 x 2.5 | 3.5 | 62° | 27 | 1.5 s |
| St-Laurent<br>et al.<br>(2015) | 12-channel | 22.5 x 22.5 | 96 x 96 | 2.35 x 2.35 | 3.5 | 62° | 27 | 1.5 s |

*Note.* All studies used a 3.0-T Siemens MAGNETOM Trio MRI scanner.

## Footnotes

1 Since the analyses performed in this study were based on those reported by St-Laurent et al., 2015, we considered the possibility that the current results may be biased by including the data from that study in the present analyses. Therefore, vividness analyses and LGA meta-analyses were also conducted on a subset of studies, excluding the data from St-Laurent et al. Although the subset yielded a weaker pattern given reduced power, regional relationships were consistent, so all analyses reported in this paper include all four studies.

2 The two studies that used image stimuli were analyzed alone to explore whether the subcortical contributions were driven by the temporal properties of video stimuli. Reliable correlations were, however, still detected.

